# Reduced resilience of brain state transitions in anti-*N*-Methyl-D-Aspartate receptor encephalitis

**DOI:** 10.1101/2022.01.24.477081

**Authors:** Nina von Schwanenflug, Juan P Ramirez-Mahaluf, Stephan Krohn, Amy Romanello, Josephine Heine, Harald Prüss, Nicolas A Crossley, Carsten Finke

## Abstract

**Objective:** Patients with anti-NMDA receptor encephalitis suffer from a severe neuropsychiatric syndrome, yet most patients show no abnormalities in routine magnetic resonance imaging. In contrast, advanced neuroimaging studies have consistently identified disrupted functional connectivity in these patients, with recent work suggesting increased volatility of functional state dynamics. Here, we investigate these network dynamics through the spatiotemporal trajectory of meta-state transitions, yielding a time-resolved account of brain state exploration in anti-NMDA receptor encephalitis.

**Methods:** Resting-state functional magnetic resonance imaging data were acquired in 73 patients with NMDAR encephalitis and 73 age- and sex-matched healthy controls. Time-resolved functional connectivity was clustered into brain meta-states, giving rise to a time-resolved transition network graph with states as nodes and transitions between brain meta-states as weighted, directed edges. Network topology, robustness, and transition cost of these transition networks were compared between groups.

**Results:** Transition networks of patients showed significantly lower local efficiency (*t* = -2.54, *p*_*FDR*_ = 0.026), lower robustness (*t* = -2.01, *p*_*FDR*_ = 0.048) and higher leap size (*t* = 2.33, *p*_*FDR*_ = 0.026) compared to controls. Furthermore, the ratio of within-to-between module transitions and state similarity was significantly lower in patients. Importantly, alterations of brain state transitions correlated with disease severity.

**Interpretation:** These findings reveal systematic alterations of transition networks in patients, suggesting that anti-NMDA receptor encephalitis is characterized by reduced stability of brain state transitions and that this reduced resilience of transition networks plays a clinically relevant role in the manifestation of the disease.

## INTRODUCTION

Anti-*N*-methyl-aspartate receptor (NMDAR) encephalitis is an immune-mediated disorder of the central nervous system caused by autoantibodies targeting the NMDA receptor and leading to a dysregulation of the glutamatergic neurotransmitter system.^1^ The disease manifests in a complex neuropsychiatric syndrome with prominent psychiatric symptoms (e.g., delusions and psychosis), but also seizures, dyskinesia, psychosis, decreased levels of consciousness, and cognitive dysfunction.^2–4^ Despite the severe disease course, only 50-70% of patients show abnormalities in standard structural magnetic resonance imaging (MRI),^3,5^ resulting in a clinico-radiological paradox. In contrast, several functional MRI studies have suggested disrupted functional connectivity (FC) in NMDAR encephalitis that is linked to disease severity, disease duration and cognitive symptoms.^2,4,6–8^

FC as measured with resting-state functional MRI (rs-fMRI) is estimated from the pairwise correlation of blood oxygen level-dependent (BOLD) activity between brain regions without the presence of an explicit task.^9^ However, traditional ‘static’ approaches obtain FC as an average across several minutes, therefore missing important information that may be derived from dynamic changes in functional connections.^10,11^ Hence, the analysis of FC has been recently refined from a time*-invariant* static account to a time-*varying* description. This methodological progress allows to unveil temporal properties of functional brain organization, such as the identification of functional states, i.e., transient connectivity patterns, and their transition trajectories. These FC dynamics are thought to reflect brain state exploration that facilitates cognition and behavior, and may vary with disease.^12–14^ Accordingly, a recent case-control study investigating FC dynamics in NMDAR encephalitis showed that patients exhibited altered state preference as well as increased transition frequencies between major connectivity patterns.^15^ However, a detailed investigation of the transition trajectory of brain states and its link to clinical symptoms is still missing. Brain state exploration - facilitated by transitions between functional states - is thought to ensure stable information representation while promoting functional integration across distant brain regions and subsystems and, if disturbed, potentially affects information integration and behavior.^13,16^ Hence, identifying mechanisms and disruptions of these transition trajectories may contribute to the understanding of the pathophysiology of NMDAR encephalitis and further neuropsychiatric diseases that are associated with NMDAR dysfunction, e.g. schizophrenia.

Graph theoretical approaches are well-suited to study the temporal architecture of state exploration.^17^ recently introduced the concept of transition networks to investigate the trajectory of traversing functional states (from hereon also referred to as meta-states following.^17^ In this concept, transition networks are represented as graphs with brain states as nodes and transitions between meta-states as directed and weighted edges. Similar to other biological systems,^18^ transition networks show properties of complex networks (i.e., heavy-tailed degree distribution, high local efficiency, and modularity) indicating an organized, cost-efficient, non-random temporal trajectory of brain states.^17^ Furthermore, transition network characteristics have been related to motor function and cognitive performance in healthy controls indicating behavioral relevance.^17^

Here, we aimed to specify alterations of the spatiotemporal trajectory of state transitions and its relation to disease severity in NMDAR encephalitis. Therefore, we constructed transition networks for a large sample of patients and age- and sex-matched healthy controls. We hypothesized that the temporal structure of state exploration in NMDAR encephalitis would show altered dynamics^15^ and weakened stability of transition networks compared to a group of healthy controls.

## MATERALS AND METHODS

### Participants

For this study, 73 patients with NMDAR encephalitis were recruited from the Department of Neurology at Charité - Universitätsmedizin Berlin. All patients fulfilled diagnostic criteria including characteristic clinical presentation and detection of IgG NMDA receptor antibodies in the cerebrospinal fluid.^3^ Patients were in the post-acute phase of their disease with a median of 2.97 years (IQR: 2.48) after disease onset. Disease severity at the time of scan and peak of disease was assessed with the modified Rankin scale (mRS). The control group consisted of 73 age- and sex-matched healthy participants without any history of neurological or psychiatric disease. Data from 49 patients and 25 controls were analyzed in a recent study by von Schwanenflug and colleagues^15^ investigating functional dynamics in NMDAR encephalitis. For the current study, patient-control matching was optimized for age and sex through a computational matching algorithm (see S1). The two groups were perfectly balanced for sex and did not differ significantly in age as tested with a Wilcoxon rank sum test (*p* = 0.61). Clinical and demographic characteristics are summarized in (Table 1). The study was approved by the ethics committee of the Charité - Universitätsklinikum Berlin and all participants gave written informed consent before participation.

**Table 1:**
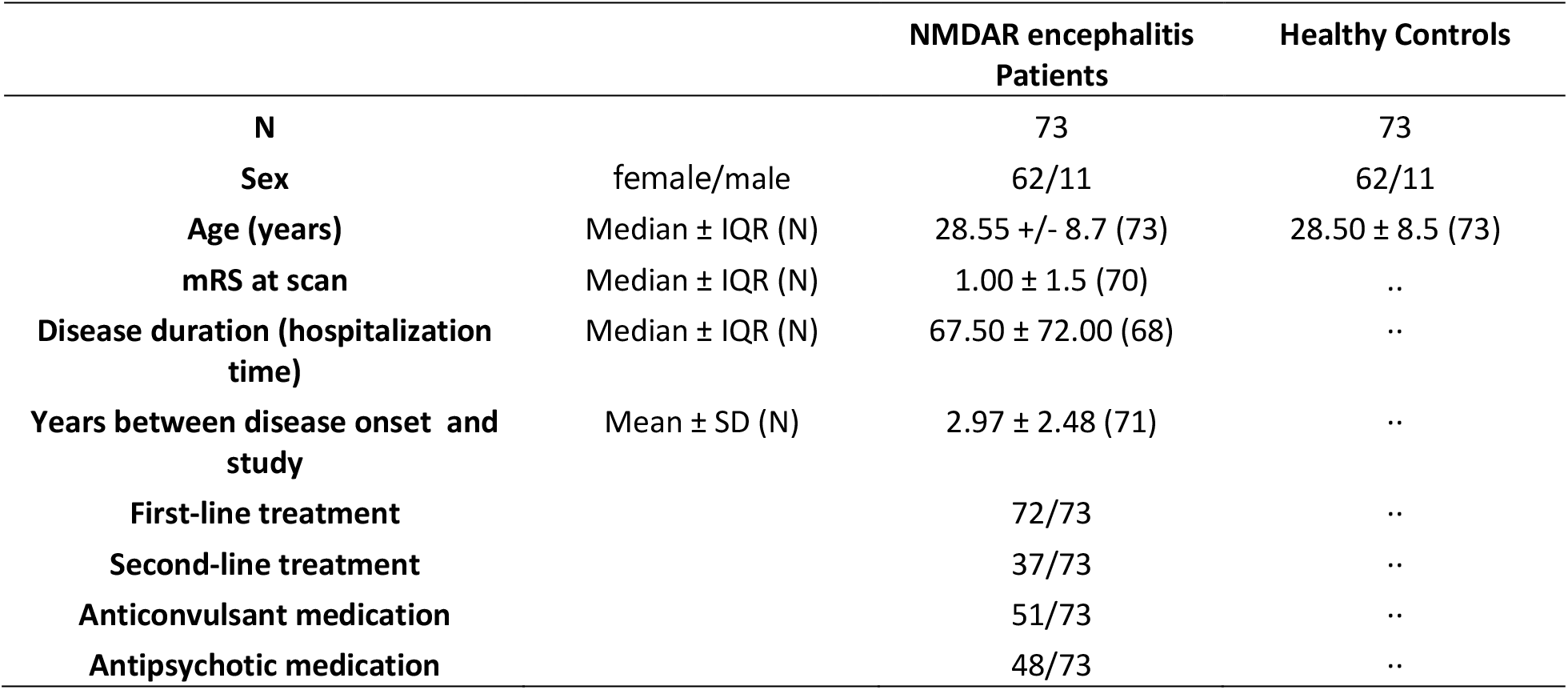
Demographic variables and clinical measures of the participants. Table lists median and interquartile range (IQR) of age, mRS at scan, mRS at peak of disease, disease duration, and time between scan and diagnosis. Treatment and medication during disease course were evaluated using a binary scale (present: ‘yes’ vs. absent: ‘no’). Disease duration = days in acute care; N = number of participants; mRS = modified Rankin Scale.

### MRI data acquisition

MRI data was collected at the Berlin Center for Advanced Neuroimaging at Charité – Universitätsmedizin Berlin using a 3T Trim Trio scanner equipped with a 20-channel head coil (Siemens, Erlangen, Germany). Resting-state functional images were acquired using an echoplanar imaging sequence (TR=2.25s, TE=30ms, 260 volumes, voxel size=3.4×3.4×3.4mm^3^). High-resolution T1-weighted structural scans were collected using a magnetization-prepared rapid gradient echo sequence (MPRAGE; voxel size=1×1×1mm^3^).

### MRI data analysis

Prior to preprocessing, framewise displacement (FD) was calculated for each participant and assessed against a mean FD cutoff of 0.50 mm.^19,20^ No participant had a mean FD greater than or equal to 0.50 mm. For preprocessing, we applied the “ICA-AROMA+2Phys”-Pipeline proposed by Parkes and colleagues^21^ to our data: the pipeline included removal of the first 4 volumes of each participant’s rs-fMRI scan, volume realignment, slice-timing correction, detrending of BOLD time-series, intensity normalization, spatial smoothing with 6mm full width at half maximum, ICA-AROMA for head motion correction to robustly remove motion-induced signal artifacts from the fMRI data,^22^ regression of white matter and CSF time-series to control for physiological fluctuations of non-neuronal origin, demeaning, and band-pass filtering to retain frequencies between 0.008 and 0.08 Hz.

### Participant-wise meta-state estimation and transition network construction

The following steps were performed with the same parameters as previously described and evaluated in Ramirez-Mahaluf and colleagues.^17^ Time series extraction was done using a whole-brain parcellation template with 638 similarly sized regions of interests (ROI).^23^ Extracted functional time series were segmented into 127 consecutive time windows of 2TRs (≙4.5 seconds), which yielded reliable results in previous work.^17^ For each window, FC was estimated between any two ROIs using Multiplicaton of Temporal Derivatives.^24^ The resulting ROI-by-ROI (638-by-638) matrices were then Pearson-correlated, resulting in a 127-by-127 similarity matrix of windows. To obtain discrete brain meta-states MATLAB-inbuilt k-means clustering was applied to the similarity matrix using 10000 maximum iterations and 2000 replicates with random initial positions. For each meta-state, all windows belonging to that state were averaged, yielding a mean ROI-by-ROI (638-by-638) connectivity matrix. To scrutinize our analyses across multiple numbers of meta-states, we extracted *k* meta-states (*k* = 35, 40, 45, 50, and 55) following the range of *k* in Ramirez-Mahaluf and colleagues^17^ for each participant separately.

Finally, transition networks were constructed for each participant and *k* number of meta-states: A transition network is a graph network, where each meta-state corresponds to a node and transitions between meta-states represent the edges of that graph. The edges are directed and weighted according to the number of transitions from meta-state *i* to meta-state *j*.^17^

Importantly, this novel approach runs k-means clustering on each individual time series to describe individual temporal trajectories of meta-state transitions. Hence this approach differs fundamentally from the definition of dynamic FC states across individuals^15^ which searches for common patterns of recurring connectivity on a group level.

### Group comparisons of transition network properties

From each of the transition networks, we derived three widely used graph theoretical measures (*modularity, local efficiency*, and *global efficiency*), two custom measures that are thought to capture the biological costs of meta-state transitions (*leap size* and *immobility*),^17^ as well as one measure that assesses the *robustness* of the network against perturbations. Note that modularity, local and global efficiency, and immobility were calculated on the transition matrix (matrix containing the number of transitions between each pair of meta-states) for each participant, while leap size was based on the distance matrix (i.e., 1-correlation for each meta-state pair). To assess robustness, we employed the *NetSwan* package available for *R* to randomly remove one node after another from the network and recalculate the size of the largest connected component.^25,26^ A more detailed description of the graph theoretical metrics is provided in (Table S1).

In addition, *transition frequency, ratio of within-to-between module meta-state similarity*, and *ratio of within-to-between module transitions* were compared between patients and controls. Here, a ‘module’ refers to a group of meta-states assigned to the same community as defined by the modularity algorithm (*community_louvain*.*m*). While transition frequency is calculated as the absolute number of transitions between different meta-states, the ratio of within-to-between meta-state similarity (ratio_*sim*_) is defined as the average correlation of meta-states within a module divided by the average correlation of meta-states between modules. Similarly, the ratio of within-to-between module transitions (ratio_*trans*_) is the absolute number of transitions within the same module divided by the absolute number of transitions between modules.

Between-group comparisons of graph theoretical measures, transition frequencies, ratio_*sim*_, and ratio_*trans*_ were assessed by comparing the area under the curve (AUC; MATLAB’s *trapz*) between patients and controls. The AUC was calculated from *k* = 35 to *k* = 55 for each metric and participant, which allowed us to derive one inference measure across all number of meta-states. For each metric separately, the AUC. was entered into a regression model controlling for head motion (FD), age, and sex as nuisance variables. Group comparisons were performed on the residuals using a permutation-based t-test and FDR-corrected using Benjamini-Hochberg.^28^

### Correlation of network properties with disease severity

Next, we investigated the relationship of transition network properties with disease severity of patients. To this end, mRS scores at the time of scanning and disease duration (days in acute care) were z-tranformed across patients and subsequently averaged, resulting in a composite z-score for each patient that reflects disease severity clinical disability. The Pearson’s correlation coefficient between dynamic network properties and disease severity was obtained and corrected for multiple comparisons.

### Functional network topology of meta-states

Lastly, each meta-state can be represented by a whole-brain FC matrix (638-by-638), in which each edge corresponds to the coupling strength between two given brain regions. Consequently, we sought to evaluate the spatial differences in functional topology of these edges across all meta-states. To this end, we quantified how much each edge differed across meta-states by computing the across-state distance (ASD), a previously defined summary measure by Krohn and colleagues^27^, which is defined as the cumulative difference across a specified state space. Here, this distance was computed for each edge across all possible meta-state comparisons given a particular value of *k*, then normalized over *k*, and finally averaged over the applied range of *k* values. In consequence, we obtain a single distance measure for each edge and participant, where a high value of ASD between any two ROIs indicates that the connectivity between these regions differs strongly between meta-states. In contrast, a low ASD indicates that the connectivity between these regions is similar across all meta-states of a transition network. Finally, the participant-specific ASD values were averaged to obtain the group-level distance matrix shown in (Fig 4). Subsequently, group differences for each edge in the distance matrix were assessed with a two-sample t-test and FDR-corrected for multiple comparisons using Benjamini-Hochberg.^28^

## RESULTS

### Group differences in network properties

Group comparisons of graph theoretical measures yielded significantly lower local efficiency (*t* = -2.41, *p*_*FDR*_ = 0.029), higher leap size (*t* = 2.18, *p*_*FDR*_ = 0.037), and lower robustness (*t* = -2.01, *p*_*FDR*_ = 0.048) of transition networks in patients compared to controls. In contrast, modularity (*t* = -1.43 *p*_*FDR*_ = 0.12), global efficiency (*t* = 1.00, *p*_*FDR*_ = 0.20), and immobility (*t* = -0.32, *p*_*FDR*_ = 0.38) of transitions networks did not differ between groups (Fig 1). (Fig S1) shows the transition networks of six exemplary participants with high and low leap size.

**Fig 1:**
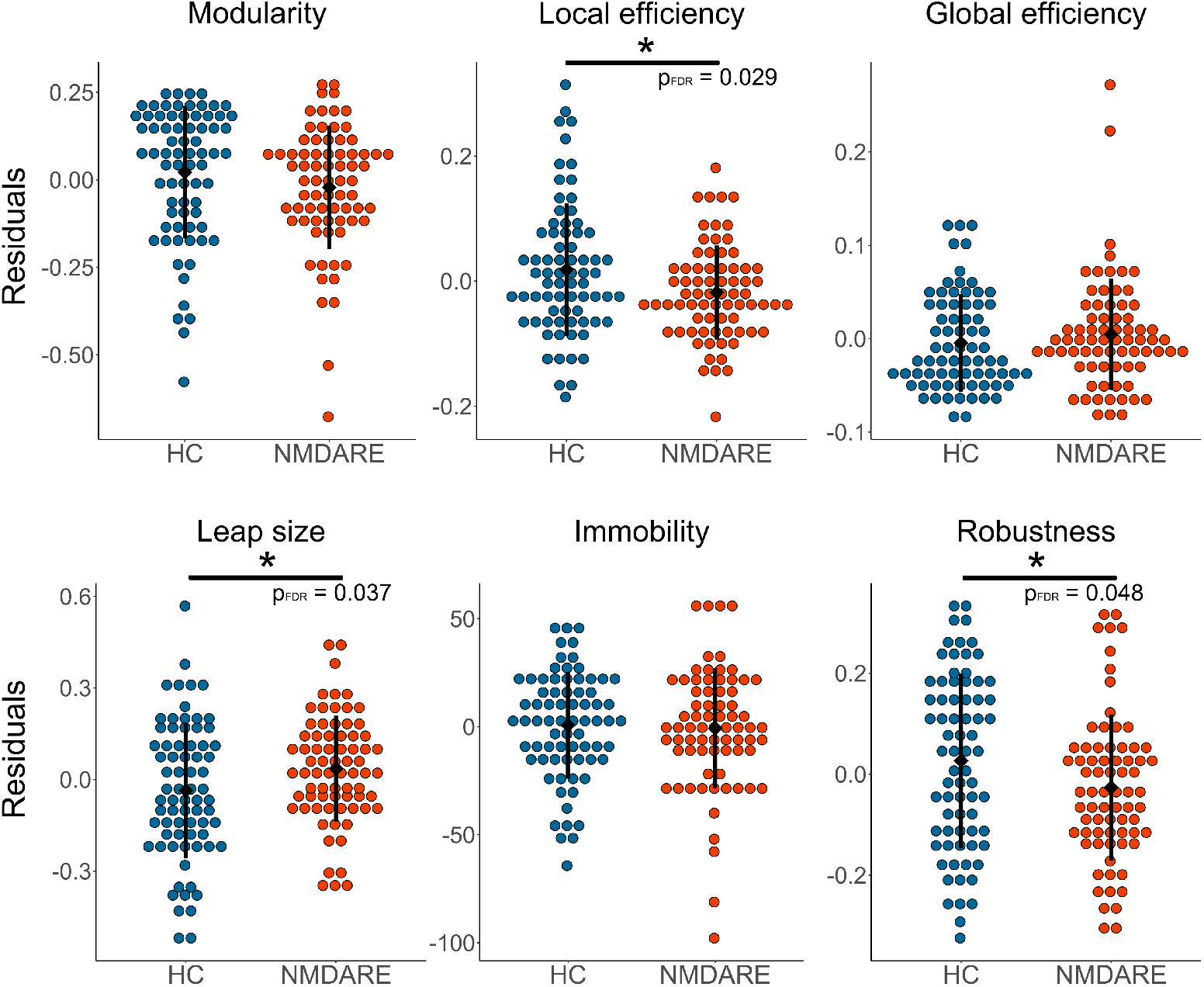
Between-group comparisons of graph theoretical measures. Colored dots represent the residuals after nuisance regression. Black dots and whiskers represent the mean and standard deviation, respectively. HC = healthy controls, NMDAR encephalitis = patients with anti-NMDA receptor encephalitis. * indicates significant difference *p*_*FDR*_ < 0.05.

Correlation of similarity and the number of transitions between two meta-states revealed that transition frequency was higher between similar meta-states for both groups and all numbers of meta-states (rho = [0.48, 0.53, 0.54, 0.54, 0.52] for the different *k* meta-states; all p < 0.001, Fig 2). Accordingly, transitions within modules were on average 3.4 times more likely than between modules with a ratio_*trans*_ (ratio of within-to-between module transitions) = [2.40, 2.94, 3.44, 3.89, 4.44] depending on the number of meta-states. This result was expected as modularity is calculated on the transition matrix. Interestingly, however, the ratio_*trans*_ was significantly lower in patients with NMDAR encephalitis compared to controls (*t* = -2.48, *p*_*FDR*_ = 0.026), while the overall number of transitions between different meta-states did not differ between groups (*t* = 0.32, *p*_*FDR*_ = 0.377). Similar to ratio_*trans*_, ratio_*sim*_ (ratio of within-to-between meta-state similarity) was on average 3.2 (ratio_*sim*_ = [2.9, 3.1, 3.2, 3.3, 3.4], for *the different k* meta-states). Again, the ratio_*sim*_ was significantly lower in patients compared to controls (*t* = -2.48, *p*_*FDR*_ = 0.026). This suggests that patients transition between topologically more different meta-states (from different modules) compared to controls, while the overall transition frequency remains unaltered.

**Fig 2:**
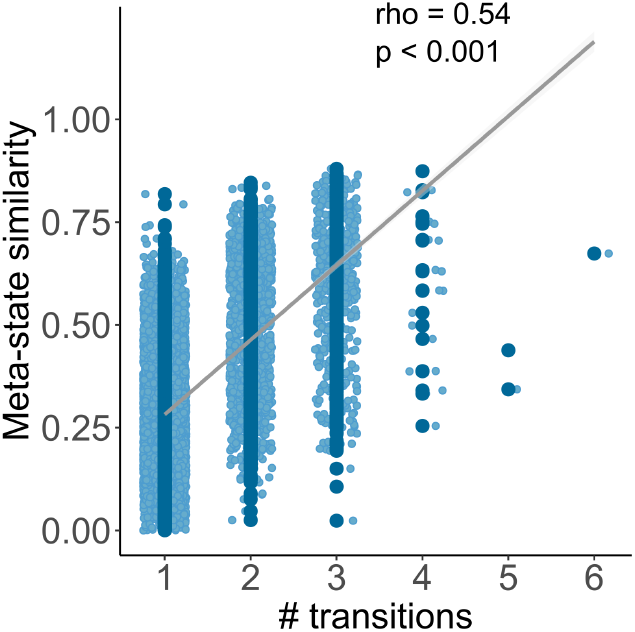
Correlation between meta-state similarity and number of transitions between them (here shown for *k* = 45, see Fig S2 for *k*’s = [35 40 50 55]). Meta-state similarity (y-axis) was estimated calculating Spearman’s ϱ. The regression line is included for visualization purposes. Number of transitions (x-axis) are the sum of transitions between any two meta-states, independent of the direction of transitions.

### Correlation of network properties with disease severity

Next, we investigated the relationship of significant graph metrics, i.e., local efficiency, leap size, and robustness, ratio_*trans*_ and ratio_*sim*_, with a composite z-score for disease severity. Higher disease severity was significantly associated with higher leap size (Pearson’s *r* = 0.37, *p*_*FDR*_ = 0.0030, Fig 3), decreased robustness (Pearson’s *r* = -0.37, *p*_*FDR*_ = 0.0030, Fig 3), lower ratio_*sim*_ (Pearson’s r = -0.40, *p*_*FDR*_ = 0.0030, Fig S3), and lower ratio_*trans*_ (Pearson’s r = -0.33, *p*_*FDR*_ = 0.0064, Fig S3), but not with local efficiency (Pearson’s r = -0.11, *p*_*FDR*_ = 0.35, Fig S3).

**Fig 3:**
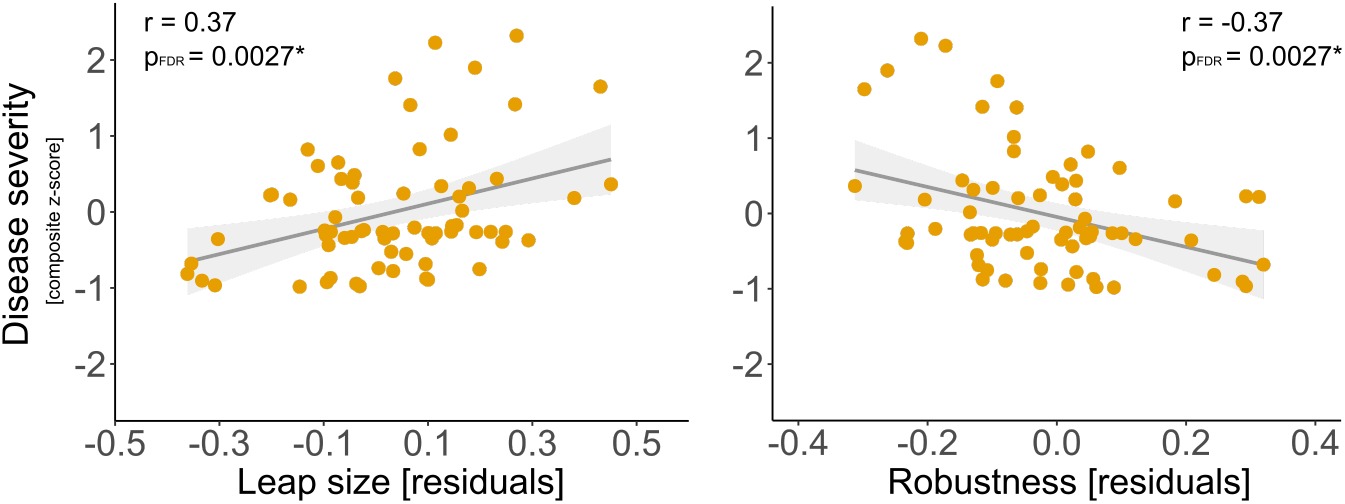
Correlation between disease severity (composite z-score) and altered network properties (residuals after nuisance regression). Correlation plots for local efficiency, ratio_*sim*_ and ratio_*trans*_ are shown in Fig S3. * indicates significant difference *p*_*FDR*_ < 0.05.

**Fig 4:**
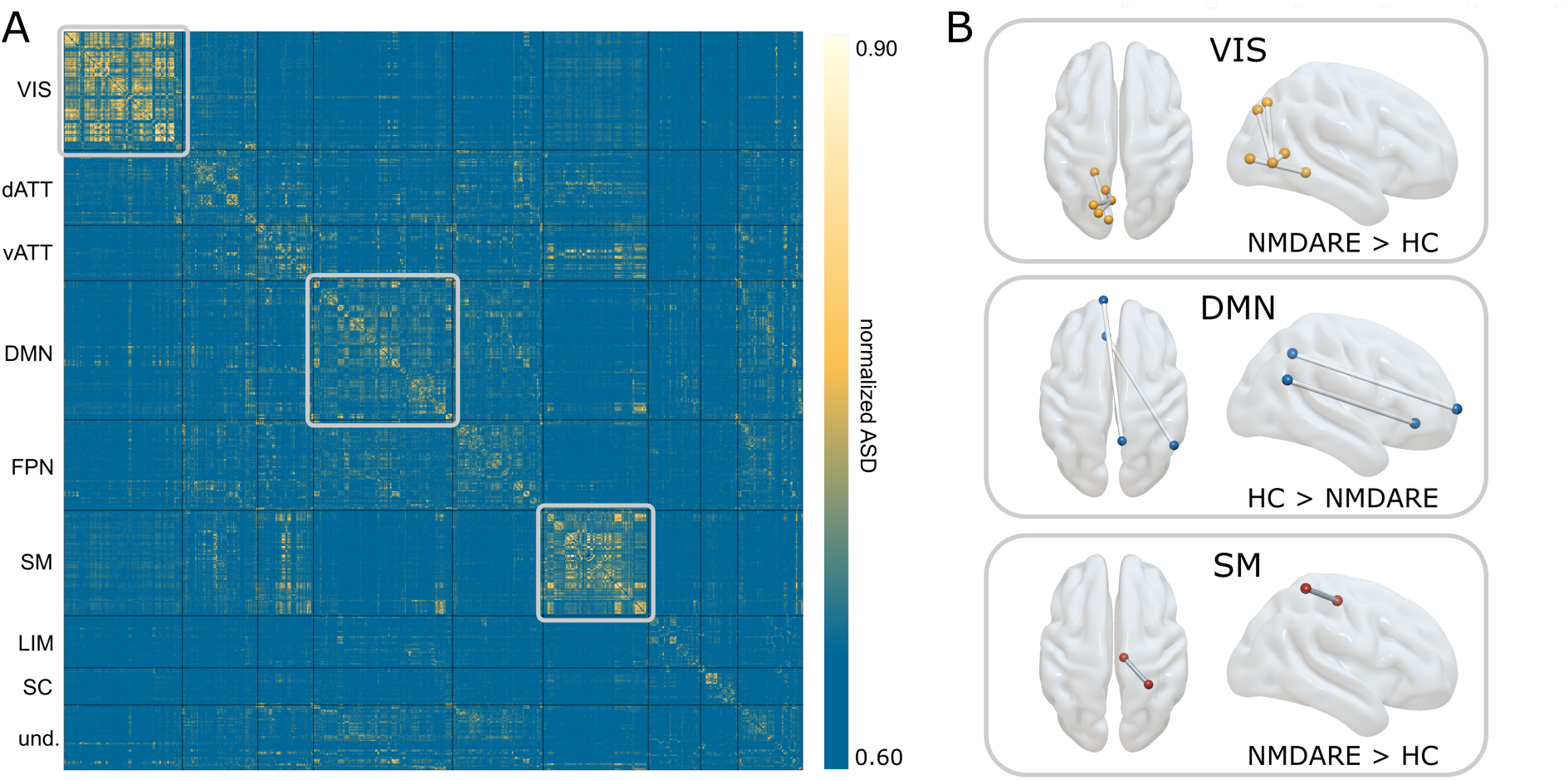
Interregional across-state distance (ASD) matrix averaged across all participants. Network assignment of regions is based on the labels proposed by Yeo et al.^44^ Subcortical regions were subsumed as a subcortical network. Brain plots show results from group-comparison within each functional network. Differences in ASD between patients and healthy controls were found for edges within the visual, default-mode, and sensorimotor network (FDR-corrected). VIS = visual network, dATT = dorsal attention network, vATT = ventral attention network, DMN = default mod network, FPN = fronto-parietal network, SM = sensorimotor network, LIM = limbic network, SC = subcortical network, und. = undefined, HC = healthy controls, NMDAR encephalitis = patients with anti-NMDA receptor encephalitis.

### Functional network topology of meta-states

The edges with the highest ASD, i.e., edges that exhibited most pronounced differences in coupling strength across meta-states, clustered predominantly in unimodal networks, namely the sensorimotor and visual network (Fig 4A). This topological pattern is highly convergent with recent findings from Krohn and colleagues.^27^

Whole-brain group comparison yielded no significant difference in ASD between groups after correction for multiple comparison. Therefore, we explored group differences in ASD within each functional resting-state network separately. This network-wise group comparison revealed significant differences between edges within the visual, default mode and sensorimotor networks (FDR-corrected, Fig 4B). Remarkably, significant edges showed higher ASD in patients within the visual and sensorimotor network, but lower ASD within the default-mode network (Table S3).

## DISCUSSION

Ongoing brain activity can be described as transient FC patterns (so-called brain states) that are visited in a structured, non-random trajectory.^17^ These brain state dynamics are thought to facilitate cognition and behavior, and may vary in disease.^12^ In this study, we employed a time-resolved analysis of brain activity to capture the spatiotemporal dynamics of brain state transitions in a large sample of patients with NMDAR encephalitis. Our results indicate reduced resilience of state transition networks in patients compared to controls. This manifests in lower local efficiency of the network (fewer transitions from or to neighboring, i.e., similar, meta-states), higher leap size (transitions between more distinct meta-states), and reduced robustness of the patients’ transition networks against random attacks. Furthermore, the ratio of within-to-between module transitions and meta-state similarity was significantly reduced in patients. Importantly, these state dynamic metrics were correlated with disease severity, highlighting the clinical relevance of our findings.

In patients with NMDAR encephalitis, autoantibodies target the NR1 subunit of the NMDA receptor causing an internalization of the receptor.^29^ While this results in a broad range of psychiatric and neurological symptoms, standard clinical MRI shows no or only minor abnormalities in most patients.^3,5^ In contrast, FC analyses were able to identify characteristic connectivity alterations: *Static* resting-state FC analyses that average connectivity across an entire scanning session showed widespread disrupted connectivity in visual, temporal, hippocampal, and mid-frontal areas associated with the severity of cognitive and psychiatric symptoms.^6,30,31^ However, given that brain activity is inherently dynamic,^32^ models that incorporate spatiotemporal features of connectivity may complement our knowledge about functional disruptions in neuropsychiatric disorders. Indeed, we recently found that *dynamic* FC showed a shift in state preference and transition probabilities in patients with NMDAR encephalitis that was associated with disease severity and disease duration.^15^ In the present study, we further expand on these dynamic FC findings and investigated alterations in the *spatiotemporal trajectory* of functional state exploration through the underlying state space. State exploration is thought to reflect the dynamic repertoire of intrinsic brain activity that is important for information integration and mental processes.^13,16,33^ Therefore, disruptions in the temporal organization of state transitions may account for clinical symptoms in disease.^12,34^ In fact, we found a characteristic spatiotemporal reorganization of the transition trajectory in patients compared to controls that was related to disease severity. This spatiotemporal reorganization - as reflected by lower local efficiency, lower robustness, and higher leap size of the transition network - may represent overly unstable transition dynamics in NMDAR encephalitis ^15^ that could be linked to deficiencies in information integration.^16,34^

At a scale of seconds to minutes, the human brain operates through continuously evolving activity that can be characterized as transient quasi-stable brain states.^10,11^ This evolution of brain activity is non-random, allowing for a systematic exploration of brain states.^17^ Analogous to the modular spatial organization of the cortex, the temporal trajectory of brain state transitions shows similar topological properties, i.e. brain states are grouped into modules of similar meta-states, with higher transition frequencies within a module than between modules (see methods and results section: ratio_*trans*_/ratio_*sim*_). This modular organization is thought to promote segmented and cost-efficient information processing, while enabling the exploration of the functional repertoire via transitions to meta-states of a different module.^14,34–37^ In line with the modular structure, transition networks in healthy controls show high local efficiency, and low global efficiency (as compared to a null model).^17^ While a high local efficiency allows for locally specialized functioning, a comparatively smaller number of connections between subsystems of a network, i.e., low global efficiency, still allows for distributed information processing across different subsystems.^37^ Moreover, a high local efficiency enhances the robustness of a system. By providing alternative pathways between two nodes (i.e., meta-states), the system compensates for potential disturbances and provides stable representation of information.^38^ Interestingly, the spatiotemporal organization of state exploration may also be directly relevant to behavior. A recent study on transition networks in a healthy population suggests that the efficiency of the network is associated with performance in cognition and motor function.^17^ Thus, state exploration may vary across diseases potentially accounting for a multitude of symptoms.^12^

Indeed, the present study highlights significant differences in the temporal architecture of transition networks between patients with NMDAR encephalitis and healthy controls. We found that patients exhibited decreases in local efficiency and robustness, and increases in leap size. Decreased local efficiency hints at unstable representation of information due to lower redundancy of the transition network, which is described in more detail in the previous paragraph. Leap size is thought to reflect metabolic cost and is measured as the magnitude of ‘jumps’ between topologically different meta-states. Eliciting state transitions is energetically costly^33,39^ and possibly increases when traversing states that show highly disparate activation profiles. This intuition is supported by our finding that (low-cost) transitions between two similar meta-states are more likely than (cost-intensive) transitions between distant meta-states. Accordingly, higher leap size in patients may indicate higher metabolic demand along with higher volatility of state transitions. Lastly, we evaluated the robustness of the transition network, which is the ability of maintaining information processing within the network before collapsing.^40^ We found that transition networks of patients with NMDAR encephalitis are less robust compared to those of controls when removing the nodes (i.e., meta-states) one by one. Together with a decreased local efficiency and higher leap size, this points to a destabilization and reduced resilience of transition networks in patients with NMDAR encephalitis, corroborating earlier findings in this patient population.^15^ This notion is furthermore supported by decreased ratios of within-to-between module transitions and within-to-between module meta-state similarity in patients. In addition, four out of five network measures - leap size, robustness, ratio_*trans*_, and ratio_*sim*_ - were correlated with a composite score of disease severity, supporting clinical relevance of our findings. Interestingly, reduced resilience of transition networks in patients with NMDAR encephalitis is supported by findings from attractor-based computational models that postulate that NMDAR dysfunction may lead to overly unstable attractors in brain activity.^41,42^ NMDAR hypofunction, as in NMDAR encephalitis and schizophrenia, may lead to a flattening of the attractors (destabilizing effect), facilitating perturbations to provoke transitions between attractors.^41,43^

Finally, our study provides evidence that a subset of regions preferentially promotes brain state transitions.^12,27^ Convergent with recent work on brain dynamics, we found that changes in connectivity across states are most pronounced in regions of the visual and sensorimotor areas, potentially following a hierarchy from unimodal to transmodal networks.^27^ Interestingly, in patients with NMDAR encephalitis, connectivity changes across meta-states within unimodal networks were even more pronounced, providing a spatial correlate of the increased temporal volatility in state transitions. Limiting state transitions to a defined number of regions initiating those transitions raises the intriguing possibility that controlled external stimulation of these particular regions could be applied to achieve a rebalancing of state dynamics.^12,33^

Some limitations of our study deserve mentioning: First, the window length of 2 TR (≙4.5 seconds) is comparatively short and may decrease the signal-to-noise ratio. However, the size of this time window has been recognized as a good trade-off between sensitivity and specificity.^24^ Furthermore, a high reliability of meta-state estimation was previously shown using a very similar window length.^17^ Second, k-means clustering is applied to each participant separately. While this approach poses inherent limitations to study between-group differences in meta-state topology, it is particularly suited to investigate individualized temporal dynamics and characteristics of the transition trajectory of functional states. Third, the applied *k* enforces the extraction of a large number of (potentially similar) meta-states for each participant. While most studies focus on 3-5 distinct major connectivity states defined on a group level, this number of states may not be sufficient to represent the full repertoire of functional configurations of the human brain. Furthermore, a small number of states limits a detailed investigation of individual transition trajectories between these states, which was the main purpose of the present study.

## Conclusion

In this study, we employed a time-resolved graph analytical framework to study the spatiotemporal trajectory of brain state transitions in patients with NMDAR encephalitis. Besides decreases in local efficiency, we observed reduced robustness of the patients’ transition networks against random attacks compared to those of healthy controls. Together with higher leap size in patients, these findings show reduced resilience of functional state transitions in patients, that is related to disease severity. Hence, our findings add to the evidence that disturbance of functional brain network dynamics plays a key role in the pathophysiology of NMDAR encephalitis.

## Supporting information

Supplements_TransitionNetwork

## AUTHOR CONTRIBUTIONS

NS and CF designed the study, interpreted the results. CF, JH, HP conceived the cohort study and data collection. NS conducted the analyses, wrote the first draft of the study and produced the figures. JRM developed the graph transition network construction and provided analysis code. SK implemented the across-state-distance and the matching algorithm. JRM, NAC and SK significantly contributed to the design and interpretation of the analysis. AR implemented the preprocessing pipeline All authors edited and revised the manuscript.

## ACKNOWLEDGEMENTS

Funded by the Deutsche Forschungsgemeinschaft (DFG, German Research Foundation, grant numbers 327654276 (SFB 1315), FI 2309/1-1, FI 2309/2-1 and PR 1274/6-1) and Deutsches Ministerium für Bildung und Forschung (BMBF, German Ministry of Education and Research, grant number 01GM1908D, CONNECT-GENERATE). HP received funding from the Helmholtz Association (HIL-A03). NS is a doctoral scholar at Cusanuswerk – Bischöfliche Studienförderung. AR is a doctoral scholar at the Berlin School of Mind and Brain, Humboldt-Universität zu Berlin. JRM and NAC were funded by the Agencia Nacional de Investigación y Desarrollo from Chile (ANID), through its programs FONDECYT postdoctorado (Ref: 3190311 to JRM), FONDECYT regular (Ref: 1200601 to NAC), and Anillo ACT192064. The funders had no influence on study design, data collection, data analyses, data interpretation, or writing.

