## Supplements_TransitionNetwork for "Reduced resilience of brain state transitions in anti-*N*-Methyl-D-Aspartate receptor encephalitis"

**Supplemental Material**

**S1:** To optimize patient-control matching for age and sex differences, we implemented a matching algorithm using the MatchIt package (version 3.0.2, <https://github.com/kosukeimai/MatchIt>) for R (version 3.6.3)(1). Propensity scores were estimated using a generalized linear model with the default logit link function. Matches were determined with greedy nearest neighbor matching, enforcing exact sex matching. To optimize the trade-off between pairwise age deviations and sample size, we applied a series of different caliper values (c=0.01 to 0.5). Calipers represent a liberality parameter defining the number of standard deviations of the distance measure within which to draw control units (<https://r.iq.harvard.edu/docs/matchit/2.4-15/Additional_Arguments_f3.html>). Accordingly, lower caliper values reflect stricter age optimization, leading to lower age deviations but also a smaller final study sample. We here chose the pivot point of the age-vs-sample plot as the optimal liberality parameter, as allowing for more pronounced age deviations in the matching after this point only yields small increases in the overall sample size while opting for stricter age matching results in a steep drop in sample size.

Finally, some of the control participants contributed more than one scan to the set of possible control matches (ie, longitudinal data for these participants were available). To ensure that a particular control participant was only ever matched once, we randomly resampled the subset of longitudinal scans and repeated the matching procedure. Over 10000 repetitions, we thus found the best matches given the constraint that any control participant was uniquely matched to one patient.

This procedure yielded a final study population of n=146, with perfect sex matching (62 females, 11 males for both groups) and excellent age matching (healthy controls: mean age 25.68 ± 12.97 years; patients: mean age 26.40 ± 14.03 years).

**Table S1:** Anticonvulsant and antipsychotic treatment information for patients at the time of the scan. 27 patients received anticonvulsants, 8 received antipsychotics, and 42 patients received no treatment at the time of the scan.

|  | **NMDAR encephalitis patients** |
| --- | --- |
| **Anticonvulsant medication** | 27/73 |
| **Antipsychotic medication** | 8/73 |
| **none** | 42/73 |

**Table S2:** Description of graph theoretical measures assessed for between-group comparisons.

| **Metric** | **Definition** | **Reference** |
| --- | --- | --- |
| **Modularity** | Global parameter, which quantifies the degree to which the transition network can be subdivided into clearly defined communities or modules with maximally possible number of within-module links and minimally possible number of between-module links. In the present work, a high modularity indicates that meta-states within a module show particularly higher transition frequencies compared to meta-states from different modules. Applies the Louvain community detection algorithm (*community_louvain.m*) from the Brain Connectivity Toolbox to the transition matrix of each participant; measure of segregation. | (2,3) |
| **Global efficiency** | The global efficiency is the average inverse shortest path length in a network. Paths are sequences of edges denoting possible routes of information flow within a network. It is a measure of integration indicating how fast (with the least possible numbers of edges necessary) information can be transferred from one node to all other nodes in the network. In a transition network, it measures the number of transitions necessary to reach one meta-state from all other meta-states in the network. Employs the function *efficiency_wei.m* from the Brain Connectivity Toolbox. | (2,4,5) |
| **Local efficiency** | The local efficiency is the global efficiency (see above) computed on a particular node (here: meta-state). It is therefore the inverse shortest path connecting all neighbours of that node and indicates how fast and robust information processing is within the vicinity of a node. In a transition network, this measure indicates how well connected neighbouring meta-states are among each other. As local efficiency is assessed for each meta-state, the local efficiency gets averaged across all meta-states in a transition network. Employs the function *efficiency_wei.m* from the Brain Connectivity Toolbox; measure of segregation. | (2,4,5) |
| **Immobility** | In the present study, immobility quantifies the average number of windows a participant remained in the same meta-state before transitioning to a different meta-state. | (6) |
| **Leap size** | Leap size is thought to reflect metabolic cost of state transitions and is measured as the magnitude of ‘jumps’ between different meta-states. It is defined as the distance between one meta-state and the next one (1 – correlation coefficient of their connectivity matrices). Leap size is the average distance between consecutive meta-states excluding periods of immobility. | (6) |
| **Robustness** | Measure of resilience against fragmentation of a network. Nodes (meta-states) of the transition network are randomly removed one by one. Each time, the size of the largest component (number of connected nodes) is calculated and plotted against the number of nodes removed. The robustness of the network is then specified as the area under the curve plotted. A high robustness of a transition network indicates that even in the absence of several meta-states, transitions among the remaining meta-states are still possible. Employs the *NetSwan* package for *R*, function *swan_combinatory*. | (9) |

**Table S3:** Network-wise group comparison of across-state-distance (ASD). Statistics were calculated on the participant’s average ASD across k’s. Table shows indices (idx) and region-of-interest (ROI) labels according to Crossley et al.(10)

|  | **Idx** | **ROI** | ***t*** | ***p_FDR_*** |
| --- | --- | --- | --- | --- |
| **Visual network** | 247-292 | Lingual_5 (left) - Superior occipital_4 (left) | -4.13 | 0.020 |
|  | 247-340 | Lingual_5 (left) - Calcarine_5 (left) | -3.68 | 0.024 |
|  | 247-354 | Lingual_5 (left) - Superior parietal_5 (left) | -3.91 | 0.020 |
|  | 250-336 | Lingual_8 (left) - Calcarine_1 (left) | -3.97 | 0.0015 |
| **Default-mode network** | 156-449 | Superior frontal_7 (left) - Precuneus_10 (right) | 4.11 | 0.026 |
|  | 298-637 | Angular_5 (right) - Anterior cingulum_5 (left) | 4.20 | 0.010 |
| **Sensorimotor network** | 392-499 | Potscentral_10 (right) – supplementary motor area_9 (right) | -4.36 | 0.0016 |

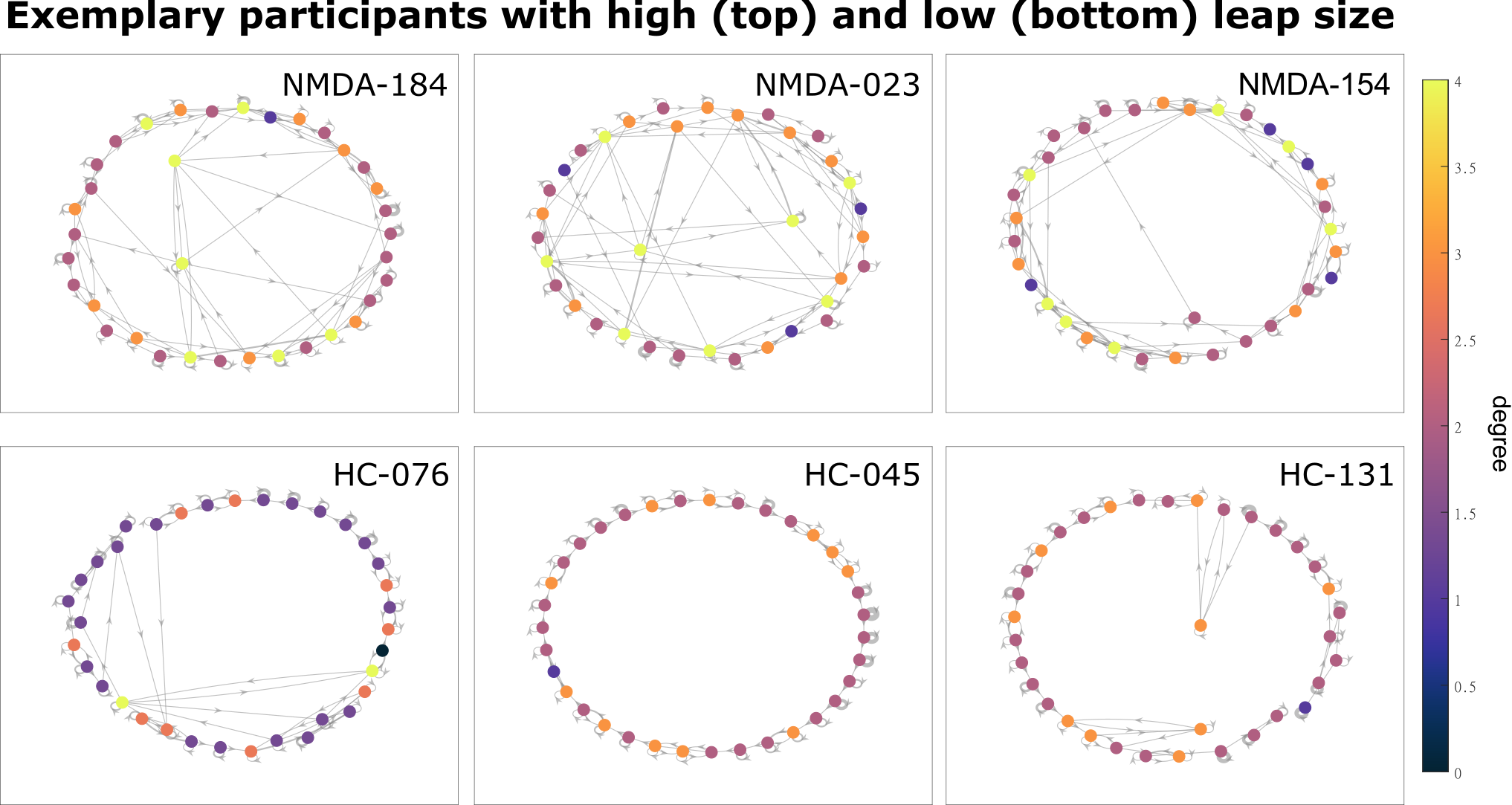

**Fig S1:** Transition networks (*k* = 35) of exemplary participants with high (top) and low leap size (bottom). Nodes are color-coded according to their degree and the distance between two nodes is proportional to their transition cost (1-correlation). The width of the edges (grey) denotes the number of transitions between the edges, while the arrows indicate the direction of transition. Self-connections represent immobility periods, i.e., no changes of meta-states between two consecutive time-windows.

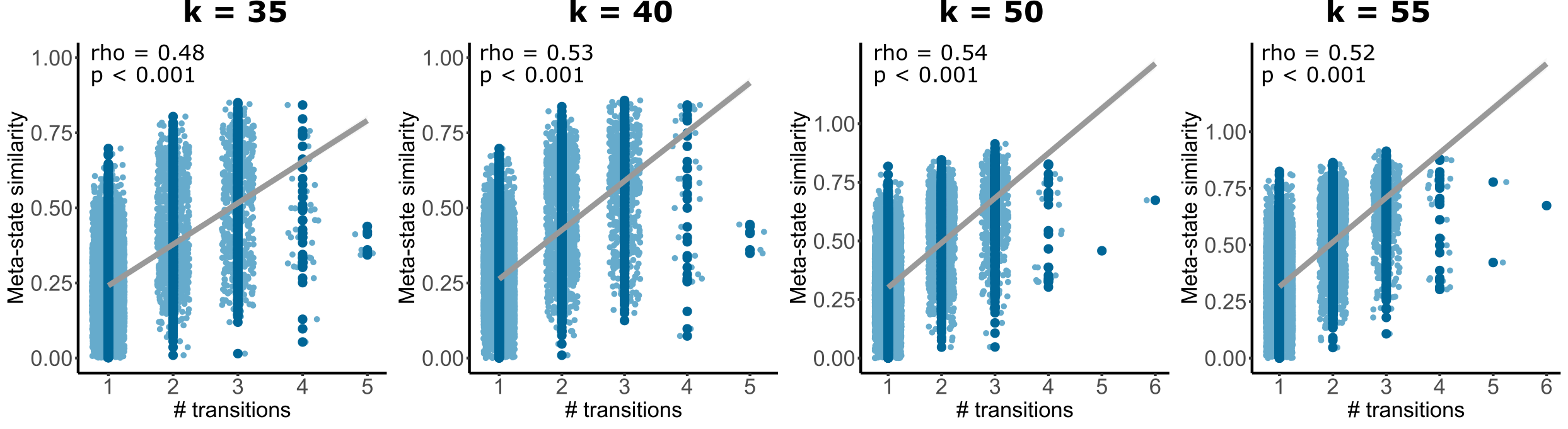

**Fig S2:** Correlation between meta-state similarity and number of transitions between them for the different number of meta-states (*k* = 45 is shown in Fig 2 in the main manuscript). Meta-state similarity (y-axis) was estimated calculating Spearman’s ϱ. The regression line is included for visualization purposes. Number of transitions (x-axis) are the sum of transitions between any two meta-states, independent of the direction of transitions.

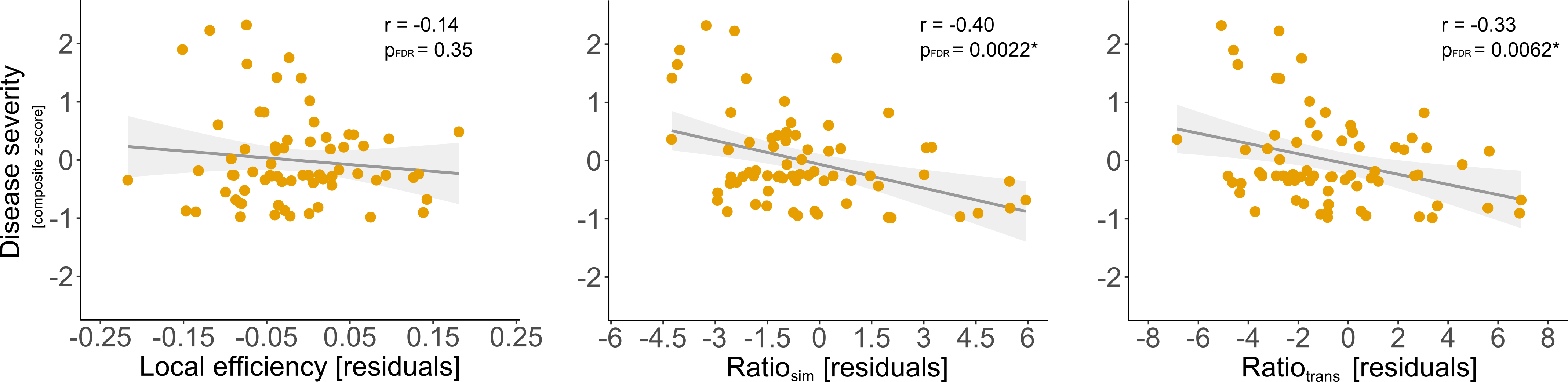

**Fig S3:** Correlation between disease severity (composite z-score) and altered network properties (residuals after nuisance regression). Correlation plots for leap size and robustness are shown in Fig 3 in the main manuscript. * indicates significant difference p_FDR_ < 0.05.
